# Wide-Field Stokes Polarimetric Microscopy for Second Harmonic Generation Imaging

**DOI:** 10.1101/2022.06.12.495834

**Authors:** Leonardo Uribe Castaño, Kamdin Mirsanaye, Lukas Kontenis, Serguei Krouglov, Margarete K. Akens, Virginijus Barzda

**Affiliations:** Department of Physics, University of Toronto, Ontario, Canada; Department of Chemical and Physical Sciences, University of Toronto Mississauga, Ontario, Canada; Laser Research Centre, Faculty of Physics, Vilnius University, Vilnius, Lithuania; Light Conversion, Vilnius, Lithuania; Department of Medical Biophysics, University of Toronto, Toronto, Ontario, Canada; Techna Institute, University Health Network, Ontario, Canada; Department of Surgery, University of Toronto, Toronto, Ontario, Canada

## Abstract

We employ wide-field second harmonic generation (SHG) microscopy together with nonlinear Stokes polarimetry for quick ultrastructural investigation of large sample areas (700 *μm* x 700 *μm*) in thin histology sections. The Stokes vector components for SHG are obtained from the polarimetric measurements with incident and outgoing linear and circular polarization states. The Stokes components are used to construct the images of polarimetric parameters and deduce the sample maps of achiral and chiral nonlinear susceptibility tensor components ratios and cylindrical axis orientation in fibrillar materials. The imaged histology sections with polarimetric wide-field SHG microscopy provide large area maps of ultrastructural information about the collagenous tissue, which can be used for rapid histology investigations.

## 1. Introduction

Polarimetric second harmonic generation (SHG) microscopy can be used for ultrastructural characterization of noncentrosymmetric materials [1–5]. It is conveniently applied for investigation of collagen structure in the histology samples [6–9]. For digital histopathology, a large area imaging is necessary, which is a limiting factor when laser scanning microscopy of SHG is employed. In contrast, the wide-field microscopy provides fast imaging of a large area due to parallel detection of the signal onto a pixel array of a camera [10, 11]. Video rate imaging can be readily performed with a wide-field nonlinear microscope [12].

The SHG intensity and its polarization sensitivity can be utilized to infer details about collagen organization in the tissue by using polarimetric SHG microscopy techniques (P-SHG) [13–15]. Full and reduced polarimetry can be applied to deduce the susceptibility tensor component values. Stokes polarimetric parameters can be employed with linear and circular incident and outgoing polarization states in reduced polarimetry measurements to deduce the achiral and chiral molecular second-order susceptibility tensor component ratios 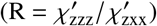 and 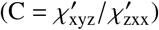, respectively, where prime signifies the susceptibility tensor of fiber projected onto the image plane with z along the projected fiber axis and y along the light propagation direction (normal to the image plane) [16–21].

In this work, a wide-field polarimetric SHG microscopy is used to obtain a number of polarimetric parameters including SHG linear dichroism (SHG_LD_), linear anisotropy of linear dichroism (LA_LD_) and circular anisotropy of linear dichroism (CA_LD_), as well as SHG circular dichroism (SHG_CD_), linear anisotropy of circular dichroism (LA_CD_) and circular anisotropy of circular dichroism (CA_CD_). The polarimetric parameters are obtained from Stokes vector components of SHG signal. The parameters then are used to calculate (without fitting) the achiral and chiral nonlinear susceptibility tensor components ratios *R* and *C* sin Δ_xyz_, and fiber orientation angles in the image plane, *δ*. Direct calculations without fitting of polarization data minimizes the post-processing time by more than an order of magnitude, which is essential for large area imaging in digital pathology. The wide-field SHG Stokes polarimetry is applied for large areas (700*μm* x 700*μm*) of rat tail tendon histology samples [21] cut at different out of section plane tilt angles, namely 0 and 45 degrees. The new wide-field Stokes polarimetric SHG imaging method can be readily applied for nonlinear digital pathology investigations.

## 2. Theoretical Considerations

### 2.1. Nonlinear Stokes-Mueller formalism for calculating outgoing Stokes vector elements

Linear and circular polarization states can be utilized for incident light as well as to probe the polarization states of generated second harmonic signal. The polarimetric parameters are deduced using double Stokes-Mueller formalism:

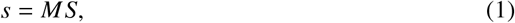

where s is the four element Stokes vector (*s*_0_, *s*_1_, *s*_2_, *s*_3_)^t^ of generated SHG signal and S is the nine element double Stokes vector for degenerate two photon polarization state of laser radiation [22]. The double Mueller matrix *M* has 4×9 elements. The double Muler matrix components are related to the nonlinear susceptibility tensor components 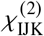 in the laboratory reference frame (see explicit relations in [23]). In turn the laboratory reference frame susceptibility components are related to the molecular susceptibility tensor components 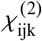 via rotation matrix with inimage-plane orientation angle *δ* and out-of-image-plane tilt angle *α* of cylindrical structures with C6 symmetry representing collagen fibers. The relations between six observable laboratory frame susceptibility components and the molecular susceptibility components are given in the Appendix. The model of chiral fibrillar aggregate is assumed where the achiral molecular susceptibilities are real, indicating that there is no phase retardance between the achiral susceptibility components, but the chiral molecular susceptibility is complex valued having a phase shift Δ_xyz_. The *R* ratio of projected susceptibility component ratio onto the image plane is defined as:

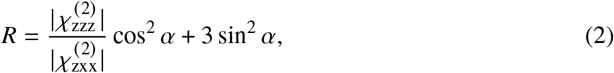

The magnitude of the chiral susceptibility component ratio, *C* is expressed as:

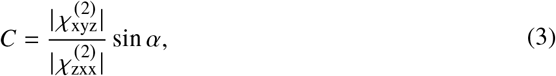

where *α* corresponds to the out-of-image-plane tilt angle of the tissue fibers. Note, that the *C* ratio is assumed to be complex valued (see the relations between laboratory and molecular frame susceptibilities outlined in the appendix) [19, 24]. We also assume that |*χ*_zxx_| = |*χ*_xxz_| [19] due to non-resonant conditions for collagen at 1030nm laser radiation wavelength.

### 2.2. Stokes polarimetric parameters with linear incident polarizations

When incident linear polarizations are employed, the SHG linear dichroism (SHG_LD_), which depends on the in-plane orientation angle *δ* of the fibrillar structures, can be readily obtained.

The SHG_LD_ is defined as:

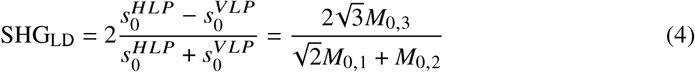

Furthermore, we can relate the SHG_LD_ to the *R* and *C* ratios, and *δ* angle as follows:

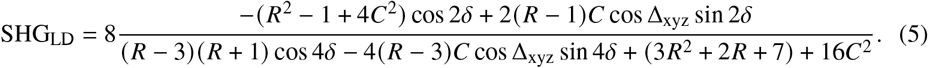

The equation shows that the signal is modulated with cos 2*δ* and cos 4*δ* functions, and depending on the sign of *C* cos Δ_xyz_, the phase of the modulation shifts slightly to the negative or positive value. The SHG_LD_ has minimum value at *δ* = 0 deg and maximum is at ±90 deg, when *C* cos Δ_xyz_ =0.

Similarly, when incident and outgoing linear polarizations are employed the linear anisotropy of linear dichroism for second harmonic LA_LD_ can be obtained. The LA_LD_ is defined as:

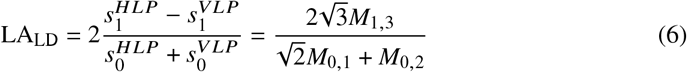

and in terms of R and C ratios, *δ* and phase shift Δ_xyz_:

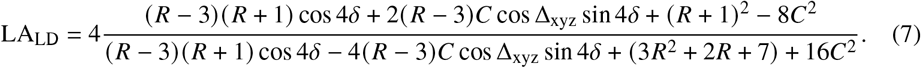

The LA_LD_ signal is modulated with cos 4*δ* dependency. There is also a significant phase shift of the modulated component, which depends on the sign of *C* cos Δ_xyz_.

If linear polarizations are used to generate SHG and the signal is subjected to a polarization state analyser (PSA) with circular polarizations, the circular anizotropy of linear dichroism CA_LD_ can be expressed as follows:

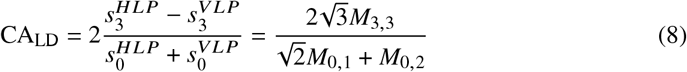

In terms of *R* and *C* ratios, *δ* and phase shift Δ_xyz_, the CA_LD_ is:

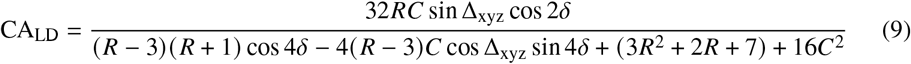

The CA_LD_ signal is modulated with periodicity of 2*δ*, and also denominator is modulated by 4*δ*. The sign and the magnitude of the CA_LD_ function depends on R and *C* sin Δ_xyz_.

### 2.3. Stokes polarimetric parameters with circular incident polarizations

Circular incident polarizations can be used with the polarization state generator (PSG) to obtain different polarimetric parameters. Total SHG signal intensity can be detected, and either circular or linear polarizations can be employed for the PSA. The polarimetric parameter of SHG circular dichroism (SHG_CD_) is defined as [21, 22, 25]:

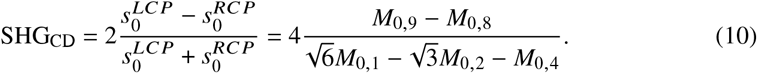

The SHG_CD_ can be expressed in terms of the R and C ratios, and the phase shift Δ_xyz_, as follows:

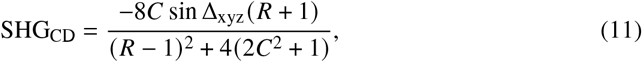

It can be seen that SHG_CD_ depends on *C* sin Δ_xyz_ and that it can flip its sign depending on the alpha orientation angle (see Eq. (3)). The SHG_CD_ also depends on the phase shift Δ_xyz_ when non-local interaction is assumed [24]. The SHG_CD_ also depends on the *R* ratio, however the sign of SHG_CD_ will not flip due to *R* dependence on the in-plane fiber angle *α* (see Eq. (2)). Hence, changing the tilt angle *α* from positive to negative values, flips the sign of SHG_CD_.

If the SHG signal generated by a circularly polarized laser beam is subjected to a PSA with linear polarizations, the linear anisotropy of circular dichroism (LA_CD_) is obtained:

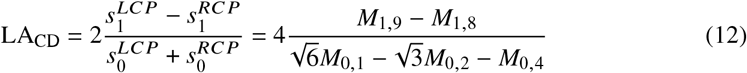

The expression in terms of *R* and *C* susceptibility ratios, *δ* and phase shift Δ_xyz_ is:

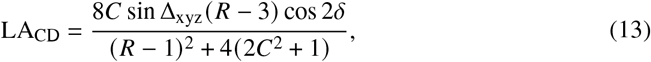

The equation of LA_CD_ shows the 2*δ* modulation of the signal. The sign of LA_CD_ directly depends on *C* sin Δ_xyz_

Analogously, the circular anisotropy of circular dichroism of SHG (CA_CD_) is defined as:

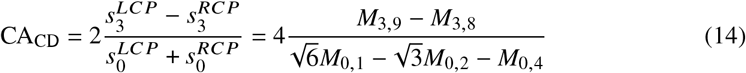

The CA_CD_ can also be expressed in terms of *R* and *C* ratios:

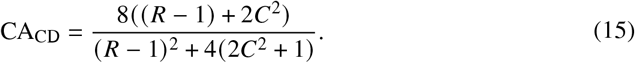

The out-of-image-plane angle *α* is explicitly included in the *R* and *C* ratios. Moreover, *C* ratio has a quadratic dependence, and when assumed to be small, it can be dropped out of the equation.

### 2.4. Ultrastructural parameters obtained from polarimetric Stokes measurements of SHG

The *R* ratio can be expressed using CA_CD_ [21]:

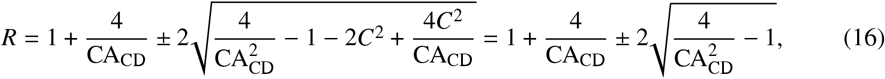

and the physically adequate range of R defines the correct sign. If *C*^2^ is assumed to be small, where (*R* – 1) ≫ 2*C*^2^, the approximate R ratio can be calculated. This equation can be used for calculating collagen *R* ratio when *C* ratio is ≤0.1.

The *C* sin Δ_xyz_ can be calculated from SHG_CD_ measurements by knowing *R* and assuming that *C*^2^ is small:

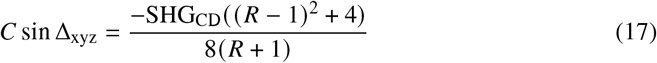

Notice that SHG_CD_ and CA_CD_ are independent of the in-plane angle of the fibrils, *δ*. In addition, we can calculate cos 2*δ* from SHG_LD_ and LA_LD_, assuming (*R* + 1) ≫ 2*C* cos Δ_xyz_:

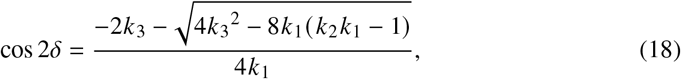

where:

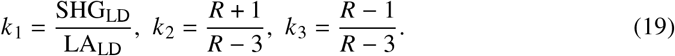

We can also obtain the in-plane orientation independent average SHG intensity when the incident circularly polarized states are employed. The intensity is expressed in terms of Stokes vector components, namely:

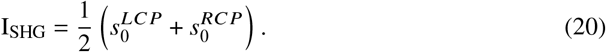

## 3. Materials and Methods

### 3.1. Wide-field polarimetric nonlinear optical microscope

A previously described wide-field polarimetric microscope was used [10, 11]. Briefly, a high power amplified femtosecond laser (Pharos, Light Conversion, Lithuania) was used for a large area wide-field illumination. The laser wavelength was 1030nm, with a pulse width of 184 fs. The laser beam was coupled to the microscope. The microscope contained a polarization state generator (PSG) comprized of a polarizing cube and a liquid crystal retarder (Thorlabs, LCC1423-A). A 75 mm focal length lens was used to focus the beam for wide-field sample illumination. The samples were placed above the focal plane. The illumination area was adjusted by translating the lens axially. The SHG signal radiated in forward direction was imaged with 20×0.5 numerical aperture (NA) air objective lens (Carl Zeiss, Germany). The polarization state analyzer (PSA) comprized of a liquid crystal retarder (Thorlabs, LCC1423-B) was placed after the objective. The SHG signal then passed through a tube lens, a linear polarizing cube, a dichroic mirror, and two filters (BG39 Shott glass filter, and 515 nm 10 nm bandwidth interference filter). The filtered SHG signal was focused onto a sCMOS camera (Hamamatsu Orca-Flash 4).

A Z-cut quartz plate and a 100 *μm* thick lithium triborate crystal attached to an electron microscope gold grid were used for the calibration of polarization setup. The SHG signal intensity had a Gaussian intensity profile along both lateral (horizontal X and vertical Z) directions in the image.

### 3.2. Sample Preparation

The tendon was harvested from a healthy rat euthanized after an unrelated study with approval of the Animal Care Committee of the University Health Network, Toronto, Canada. The tendon was formalin fixed and cut along the tendon axis (longitudinal cut) and at *α* = 45° (oblique cut). The samples were embedded in paraffin, cut into 5-*μm* thick sections and stained with hematoxylin and eosin (H&E).

## 4. Results and Discussions

### 4.1. Wide-field polarimetric SHG imaging of H&E stained rat tail tendon

Large area polarimetric imaging of H&E stained histology tissue sections is achieved without substantial fluorescence bleaching and SHG signal increase using 2 mJ/cm^2^ pulse energy density and 366 Hz pulse repetition rate [11]. A 1 second frame integration time was used to achieve the images with the signal to noise ratio of 3 (for the lowest intensity images at certain PSG and PSA configurations). The 16 different combinations of incident and outgoing polarization states were used with one retarder element PSG and PSA. One retarder PSG and PSA setup is easier to calibrate and provides quick and robust polarization measurements. The intensity images of the same sample area at various polarization states are shown in Fig. 1. The polarization states of PSG are given in the columns and polarization states of PSA are given in the rows of Fig. 1. Different areas of the sample respond differently to various incoming and outgoing polarization states. A large effect can be seen involving different circular states for PSG and PSA. The SHG images at different polarization states are used to calculate the resulting Stokes vector components maps and then to calculate the maps of polarimetric parameters.

**Fig. 1.**
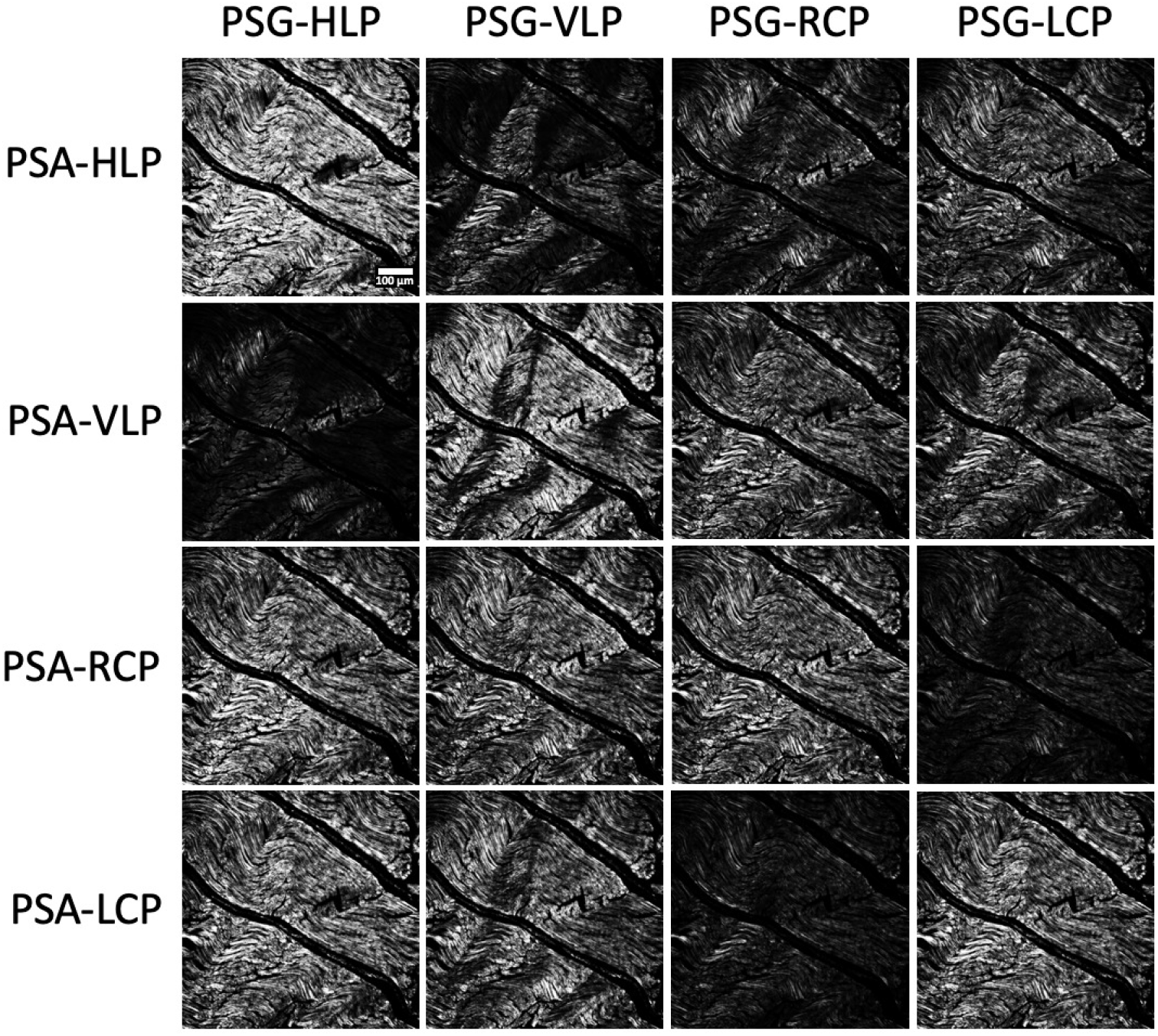
An example of longitudinally cut rat tail tendon SHG images at 16 different combinations of polarization states, measured for the polarimetric parameters calculation. The horizontally and vertically linearly polarized (HLP and VLP, respectively) and right and left circularly polarized (RCP and LCP, respectively) states were used for PSG and PSA.

### 4.2. Polarimetric parameters at different cut angles for the rat tail tendon

The maps of polarimetric parameters of SHG_LD_, LA_LD_, and CA_LD_, as well as SHG_CD_, LA_CD_,and CA_CD_ are presented in Fig. 2. The maps are calculated from the SHG Stokes vector elements using Eqs. (4), (6), (8), (10), (12), (14), respectively. Stokes vector elements are calculated from the intensity images at different PSG and PSA states (see Fig.1). The images of polarimetric parameters are calculated directly from the measured intensity images without assuming any susceptibility tensor symmetries. Therefore, the polarimetric images are obtained quickly from the raw data without any fitting requirement. For interpretation of the images of polarimetric parameters, it is informative to assume C6 symmetry with a complex valued chiral susceptibility component and use equations (5), (7), and (9) for SHG_LD_, LA_LD_ and CA_LD_, respectively, and equations (11), (13), and (15) for SHG_CD_, LA_CD_ and CA_CD_, respectively.

**Fig. 2.**
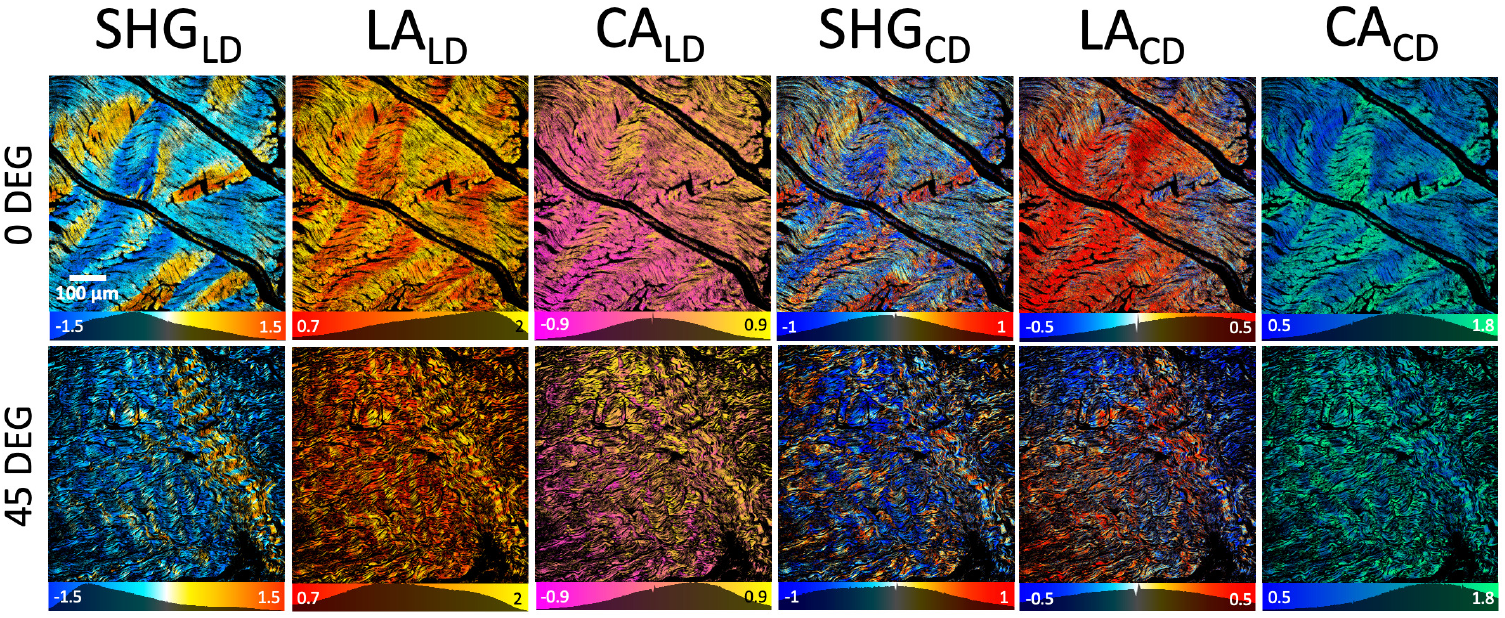
Polarimetric parameters for collagenous tissue in rat tail tendon. First row corresponds to a longitudinal cut with *α* close to 0 angle. The second row corresponds to a 45-degree cut of rat tail tendon. Images of each polarimetric parameter are presented in separate columns with labels on the column top.

The images of SHG_LD_ highlight the in-image-plane fiber orientation (Fig 2 first column). The SHG_LD_ has 2*δ* modulation of the values that reveal fiber orientation ranging from vertical at *δ*=0 degrees to horizontal at *δ*=±90 degrees. The deduced orientation angle is slightly modified compared to the actual fiber orientation due to presence of complex valued chiral susceptibility ratio (Eq. 5). For a well aligned structure of tendon cut longitudinally, the SHG_LD_ values follow closely the orientation of collagen fibers. For the oblique cut, the fiber orientations are not clearly visible, but blue pixel color indicates dominant horizontal orientation, and only a few strands are oriented towards vertical direction as indicated by the red color of the pixels (Fig 2, lower row, first panel).

The LA_LD_ has 4*δ* modulation and carries fiber orientation information. The LA_LD_ value distribution is flipped compared to SHG_LD_, and the values are positive due to addition of constant term in Eq. 7. The SHG_LD_ and LA_LD_ are used to extract the fiber orientation cos 2*δ*, as will be shown in the next section.

The CA_LD_, parameter has 2*δ* modulation, therefore it is sensitive to fiber orientation. However, CA_LD_ has strong dependence on C ratio and R ratio, and, in turn, on the tilt angle *α* out of the image plane. Therefore, the values of CA_LD_ have to be interpreted with caution, but this parameter can be employed in combination with other polarimetric parameters to extract ultrastructural information.

Polarimetric parameters with incident circular polarization states provide in-image-plane orientation free assessment of the fibers. The forth column of Fig. 2 shows SHG_CD_, which depends on the C ratio. The Δ_xyz_ influences the chiral susceptibility component that has complex values with respect to the achiral susceptibility components that are real valued [19]. C ration depends on the out-of-image-plane tilt angle *α* of the fibrils. The waviness of the collagen out and again into the image plane for the tendon longitudinal cut is nicely reflected in the SHG_CD_ image (Fig 2, first row). The SHG_CD_ is very informative at the oblique angle cut (Fig. 2 second row). The distribution of the SHG_CD_ values becomes much broader for oblique cut compared to the longitudinal cut due to *α* dependence (Eq. 3). The oblique cut image reveals islands of collagen oriented with opposite polarity, feature observed previously [18, 19]. SHG_CD_ is a very robust and quick measurement to assess collagen polarity, and can be performed with a dual or a single shot polarimetric measurement [21].

The LA_CD_ polarimetric parameter images show in-image-plane fiber orientation dependence 2*δ* (Fig. 2 fifth column). In addition LA_CD_ has strong C ratio and R ratio dependence (Eq. 13). Therefore, the interpretation of LA_CD_ images is complex due to simultaneous influence of several factors. The numerators in expressions of CA_LD_ and LA_CD_ (Eq. 9 and Eq. 13) have the same dependence on 2*δ* and C ratio, and similar dependence on R ratio, therefore the images of CA_LD_ and LA_CD_ reveal similar features (compare Fig 2, third and fifth column).

The CA_CD_ parameter is independent of the in-image-plane fiber orientation and depends on the R and *C*^2^. The CA_CD_ parameter is particularly well suited for calculating R ratio (see Eq. 16), as will be shown in the next section. CA_CD_ peak value of the distribution for longitudinal cut tendon is smaller than for the oblique cut, demonstrating the dependence of CA_CD_ on tilt angle *α*, according to Eq. 2.

The polarimetric parameters provide quick assessment of the tissue collagen ultrastructure directly from the measurements without fitting of the imaged data. The polarimetric parameters can be further employed to calculate the ultrastructure parameters.

### 4.3. Ultrastructure parameters at different cut angles for the rat tail tendon

The maps of ultrastructure parameters (Fig. 3) can be calculated from the polarimetric parameters using Eqs. (16), (17), (18), (20). The two rows of images of the ultrastructure parameters are given in Fig. 3 for comparison of longitudinal cut and oblique cut ~ 45°of the tendon. *R* ratio is calculated from CA_CD_ using Eq. (16). It slightly varies along the length of longitudinally cut tendon reflecting the out of image plane waviness of the structure. *R* ratio increases for the oblique angle cut (see *R* ratio image histograms in Fig. 3) due to *α* dependence in accordance with Eq. (2). The large area imaging of *R* ratio, which reflects the ultrastructure of collagen, can be obtained quickly by imaging with wide-field Stokes polarimetric microscopy and calculating the values without data fitting.

**Fig. 3.**
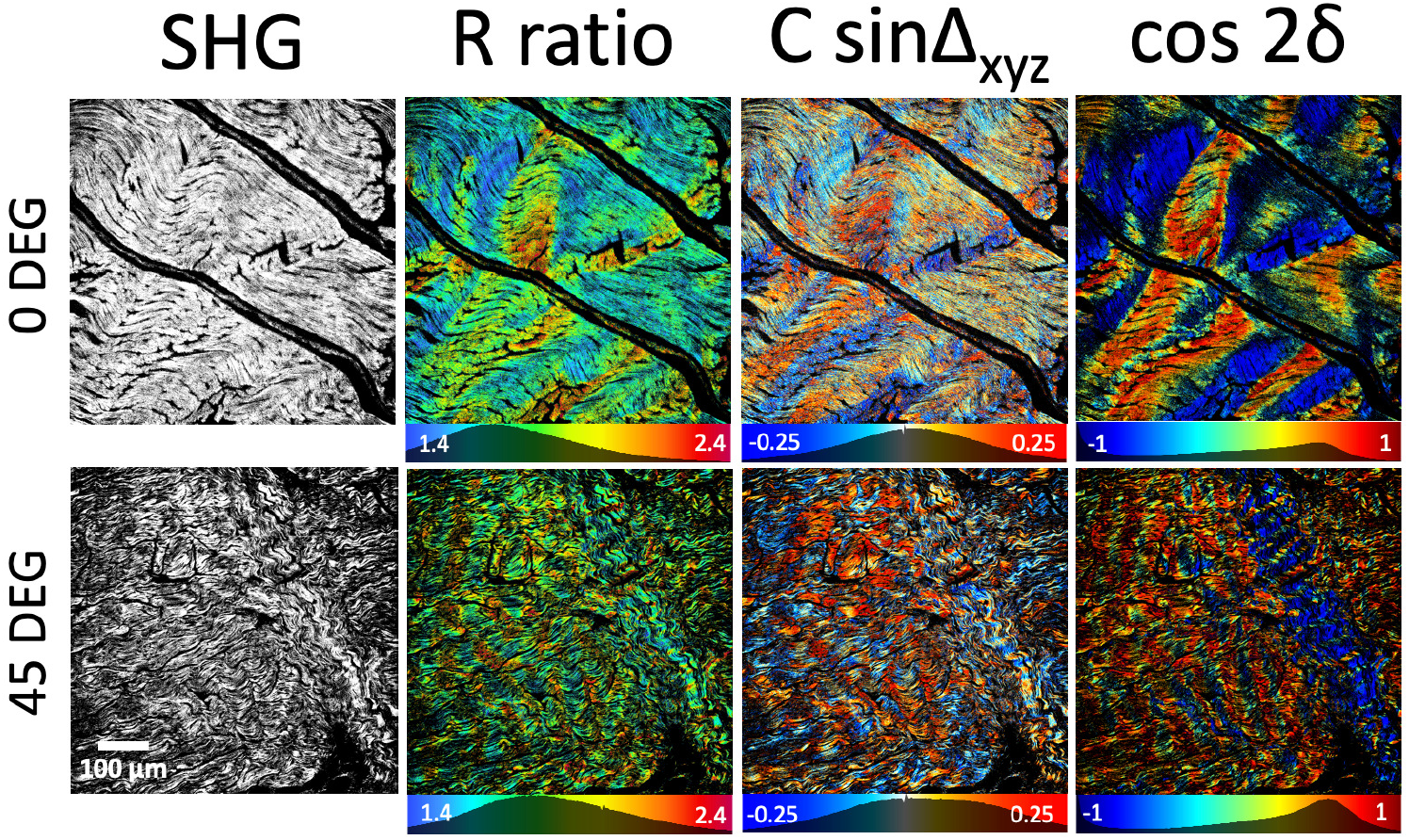
Maps of SHG intensity and ultrastructure parameters *R*, *C* sin Δ_xyz_ and orientation cos 2*δ* (respective columns) for collagen tissue in rat tail tendon. The upper row corresponds to a longitudinal cut with *α* close to 0, and the lower row correspond to a 45-degree cut of rat tail tendon. The ultrastructure parameters are calculated from polarimetric parameters shown in Fig 2.

The *C* sin Δ_xyz_ map (Fig. 3) reflects the out of image plane orientation of collagen fibers, similarly to SHG_CD_ shown in Fig. 2. This is evident due to the close relation expressed in Eq. (17). The longitudinal map of *C* sin Δ_xyz_ (Fig. 3) shows fiber tilt angle deviations from the image plane reflecting waviness of collagen fibers in the tendon. The oblique cut shows blue and red areas reflecting different polarity of collagen strands in the tendon. The *C* sin Δ_xyz_ value histogram for oblique cut is broader compared to the longitudinal cut due to larger *α* angle values.

The cos 2*δ* map shows a very clear representation of in-image-plane orientation of the collagen fibers. For the longitudinal cut (upper row, Fig. 3) the fiber in-image-plane orientation follows a wave pattern; for the oblique cut (lower row, Fig. 3), the collagenous tissue exhibits homogeneous directionality, with the exception of one cluster of fibers that are organized perpendicularly.

The reduced polarimetric measurements of tissue sections with wide-field microscopy enables large area ultrastructural assessment of collagen, which extends the utility of SHG imaging for various applications including cancer diagnostics [26, 27] and monitoring live dynamics, for example, in contracting muscle fibers [12].

## 5. Conclusions

The wide-field polarimetric SHG microscopy imaging can be used for high-throughput comparative assessment of collagen organization in histopathology samples. The images of Stokes parameters can be obtained from polarimetric measurements with both linear and circular incident and outgoing polarizations. The Stokes parameters are used to calculate polarimetric parameters, namely SHG_LD_, LA_LD_, and CA_LD_, as well as SHG_CD_, LA_CD_, and CA_CD_. In turn, the ultrastructure parameters *R*, *C* sin Δ_xyz_ and cos 2*δ* can be calculated from the polarimetric parameters without fitting. The ultrastructure parameters, as well as polarimetric parameters provide with a rapid way of characterizing ultrastructure of collagen in the tissue for histopathology investigations.

## 6. Appendix

The relations between six observable laboratory frame susceptibility components and the molecular susceptibility components are as follows:

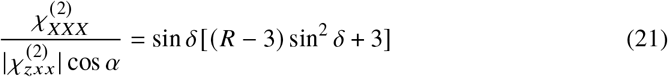

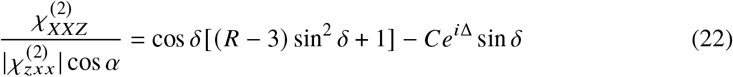

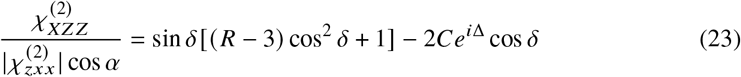

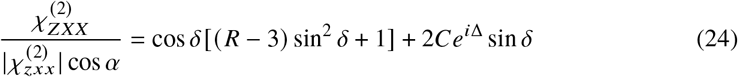

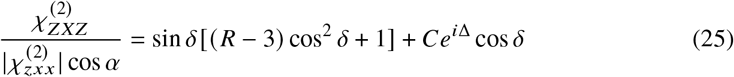

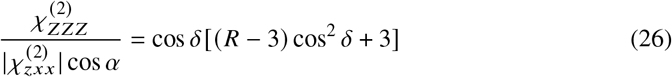

## Acknowledgements

Authors would like to acknowledge Light Conversion Inc. for landing the laser for experiments.

## Funding

The work was supported by Natural Sciences and Engineering Research Council of Canada (NSERC) (RGPIN-2017-06923, DGDND-2017-00099), and the European Regional Development Fund (project No 01.2.2.-LMT-K-718-02-0016) under grant agreement with the Research Council of Lithuania (LMTLT).

## Disclosures

The authors declare no competing financial interests.

## Notes

### Competing Interest Statement

The authors have declared no competing interest.

